# Multi-allelic *APRR2* Gene is Associated with Fruit Pigment Accumulation in Melon and Watermelon

**DOI:** 10.1101/542282

**Authors:** Elad Oren, Galil Tzuri, Lea Vexler, Asaf Dafna, Ayala Meir, Uzi Saar, Adi Faigenboim, Merav Kenigswald, Vitaly Portnoy, Arthur A Schaffer, Amnon Levi, Edward S. Buckler, Nurit Katzir, Joseph Burger, Yaakov Tadmor, Amit Gur

## Abstract

Color and pigment content are important aspects of fruit quality and consumer acceptance of cucurbit crops. Here, we describe the independent mapping and cloning of a common causative *APRR2* gene regulating pigment accumulation in melon and watermelon. We initially show that the *APRR2* transcription factor is causative for the qualitative difference between dark and light green rind in both crops. Further analyses establish the link between sequence or expression level variations in the *CmAPRR2* gene and pigments content in the rind and flesh of mature melon fruits. GWAS of young fruit rind color in a panel composed of 177 diverse melon accessions did not result in any significant association, leading to an earlier assumption that multiple genes are involved in shaping the overall phenotypic variation at this trait. Through resequencing of 25 representative accessions and allelism tests between light rind accessions, we show that multiple independent SNPs in the *CmAPRR2* gene are causative for the light rind phenotype. The multi-haplotypic nature of this gene explain the lack of detection power obtained through GBS-based GWAS and confirm the pivotal role of this gene in shaping fruit color variation in melon. This study demonstrates the power of combining bi- and multi-allelic designs with deep sequencing, to resolve lack of power due to high haplotypic diversity and low allele frequencies. Due to its central role and broad effect on pigment accumulation in fruits, the *APRR2* gene is an attractive target for carotenoids bio-fortification of cucurbit crops.

## Introduction

Flesh and rind pigmentation are key components affecting the nutritional value and consumer preference of the major cucurbits crops, melon and watermelon. Both crops exhibit extreme diversity in fruit traits, including size, shape, color, texture, aroma and sugar content (Burger *et al.*, 2006b; Wehner, 2008). Regulation of rind color in cucurbits initiates early in fruit development and is expressed as green color intensity at the young fruit stage, reflecting chlorophyll concentrations (Tadmor *et al.*, 2010). Most watermelons remain green at maturity with chlorophyll being their main rind pigment, and therefore rind color variation in watermelon is mostly expressed as green pigment intensity in uniform or striped patterns (Gusmini and Wehner, 2005). Conversely, melon rind color transforms during development, leading to extensive variation in mature fruit pigment profiles that include different combinations of carotenoids, flavonoids and chlorophylls (Burger *et al.*, 2010; Tadmor *et al.*, 2010). The genetic basis of this variation is only partly resolved. Several external fruit color QTLs have been mapped in populations derived from a cross between *Piel de Sapo* line and *PI16375* (Monforte *et al.*, 2004). It has been previously reported that the mature yellow rind color of yellow casaba melon accessions (*C. melo*, var inodorous) is caused by the accumulation of naringenin chalcone, a yellow flavonoid pigment (Tadmor *et al.*, 2010). A Kelch domain-containing F-box protein coding gene (*CmKFB*) on chromosome 10 was identified as causative for the naringenin chalcone accumulation in melon fruit rind (Feder *et al.*, 2015). While it is logical to assume that young and mature fruit color intensity are correlated, and that common genetic factors may be involved, thus far, such genes have not been reported in melon.

Three major flesh color categories are defined in melon: green, white and orange, with β-carotene and chlorophyll being the predominant pigments of the orange and green phenotypes, respectively (Burger *et al.*, 2010). The major locus qualitatively differentiating between orange and non-orange flesh is green flesh (*gf)*, located on chromosome 9 (Cuevas *et al.*, 2009). *gf* was recently shown to be the *CmOr* gene, which governs carotenoids accumulation and orange flesh color (Tzuri *et al.*, 2015). A second qualitative flesh color locus, white flesh *(wf)*, which is associated with the difference between white and green flesh, has been previously described and mapped to chromosome 8 (Clayberg, 1992; Monforte *et al.*, 2004; Cuevas *et al.*, 2009). Another layer of quantitative variation in flesh pigment content and color intensity exists within those color classes as defined by several QTL mapping studies (Monforte *et al.*, 2004; Cuevas *et al.*, 2008, 2009; Paris *et al.*, 2008; Harel-Beja *et al.*, 2010; Diaz *et al.*, 2011). These include the recent fine-mapping to a candidate causative gene level of flesh carotenoids QTL using a recombinant inbred lines (RILs) population (Galpaz *et al.*, 2018). Thus far, however, causative genes governing this quantitative variation have not been shown.

In recent years, a few transcription factors involved in regulation and synchronization of chlorophyll and carotenoids accumulation were identified in plants. Among these, Golden2-like (GLK2) transcription factors, which regulate chloroplast development (Chen *et al.*, 2016). Allelic or expression variation in the GLK2 gene were shown to be associated with levels of chlorophyll and carotenoids in Arabidopsis, tomato and pepper (Waters *et al.*, 2008, 2009; powell *et al.*, 2012; Brand *et al.*, 2014). A related but distinct transcription factor, the *ARABIDOPSIS PSEUDO-RESPONSE REGULATOR2-LIKE* gene (*APRR2*) with phenotypic effects comparable to those of GLK2, was identified and shown to regulate pigment accumulation in tomato and pepper (Pan *et al.*, 2013) and over expression of the *APRR2* gene in tomato increased the number of plastids and the color intensity. Recently, the *APRR2* gene was also shown to be causative of the white immature rind color (*w*) in cucumber (*Cucumis sativus*) (Liu *et al.*, 2016), a close relative of melon (*Cucumis melo*). The white rind phenotype of immature cucumbers in this study was shown to be associated with reduced chloroplast number and chlorophyll content.

In the current study, we used bi-parental populations to map and identify the *APRR2* gene as a common causative regulator of pigment accumulation in both melon and watermelon. We show that the effect of this transcription factor on pigment accumulation is initially observed in the rind of young fruits (chlorophylls) and extends to rind and flesh of mature melon fruits (chlorophylls and carotenoids). Through further analysis of wider genetic variation in melon, we revealed a unique multi-allelic pattern that inhibited our ability to detect a significant signal through GBS-based GWAS. By zooming in on this allelic series, we confirmed the central role of this gene in shaping the color variation of young fruit rind across melon diversity.

## Materials and methods

### Plant materials and field trials

The germplasm used in this study included four sets: (1) TAD×DUL RILs (F_7_) – bi-parental segregating population derived from the cross of the dark rind line, ‘Dulce’ (DUL; *C. melo* var. *reticulatus*) with the light rind line, ‘Tam Dew’ (TAD; *C. melo* var. *inodorous*) (Tzuri *et al.*, 2015). One hundred and sixty-four F_7_ recombinant inbred lines were developed through single-seed-descent. All RILs, F_1_ and the parental lines were grown in a randomized block design (RCBD) in an open field at Newe Ya’ar Research Center, in the spring–summer seasons of 2016 and 2017. Each line was represented by two replicates of five plants per plot. (2) NA×DUL F_3_ and F_3:4_ – An F3 population from this cross was grown in two repetitions in a greenhouse at Beit Elazari, Israel in 2013 as previously described (Ríos *et al.*, 2017). This population is derived from the cross between the light rind line ‘Noy-Amid’ (NA; *C. melo* var. *inodorous*) and the common dark rind parent, DUL. One hundred and fourteen F_3:4_ families from F_3_ genotyped plants alongside the parental lines and their F_1_ were grown in RCBD in an open-field trial at Newe Ya’ar Research Center in the spring–summer season of 2017 in two replicates of six plants per plot. (3) Melo180 GWAS panel - a Newe-Ya’ar melon collection used in this study comprised 177 diverse accessions that represent the two melon subspecies (ssp. *agrestis* and ssp. *melo*) and 11 taxonomic groups. Each accession was represented by three plots of five plants each in a randomized block design (RCBD) in the open field at Newe-Ya’ar in summer 2015 (Gur *et al.*, 2017). (4) NY0016 x EMB F_2:3_ - for mapping the light rind trait in watermelon, the light rind inbred accession NY0016 was crossed with the canary yellow accession Early Moon Beam (EMB) to produce 87 F_2:3_ families (Branham *et al.*, 2017). During the summer of 2016 and 2017, ten plants per F_2:3_ family and two plots of ten plants from the parents and F_1_ were sown in the open field at Newe-Ya’ar. All the populations used in this study were grown under standard horticultural conditions open fields at Newe Ya’ar Research Center, northern Israel (32°43′05.4″N 35°10′47.7″E), soil type was grumusol, and the plants were drip-irrigated and drip-fertilized.

### Fruit color phenotyping

In the melon populations, fruit images were taken with a digital camera on developing fruits of all accessions throughout the season, from anthesis to harvest. Rind color of young fruits was scored in the field at 10 to 15 days after anthesis and confirmed based on fruit images from the same developmental stage. Mature rind and flesh color were measured on ripe fruits, which were harvested based on abscission in climacteric fruits, or days after anthesis, rind color and TSS in non-climacteric fruits. Five mature fruits per plot were photographed externally, then cut along the longitudinal section and scanned for internal imaging, using a standard document scanner (Canon, Lide120) as described previously (Gur *et al.*, 2017). Scanned images were analyzed using the Tomato Analyzer software (Rodríguez *et al.*, 2010) for color (L, A, B, Chroma and Hue) and morphological features. Rind and flesh tissues were sampled into 50 ml tubes from at least three fruits per plot, immediately frozen in liquid nitrogen and then stored at −80°C for further analyses. For the watermelon mapping experiment (NY0016×EMB F_2:3_), ten F_3_ individuals per F_2:3_ family were harvested at maturity (∼70 days post sowing), imaged and phenotyped for rind color as above.

### Carotenoids and chlorophyll quantification

Carotenoids were extracted from 0.5 mg ground tissue samples in a mixture of hexane:acetone:ethanol (2:1:1, v/v/v) as described previously (Tadmor *et al.*, 2005), and separated using a Waters 2695 HPLC apparatus equipped with a Waters 996 PDA detector (Milford, MA). Carotenoids were identified by their characteristic absorption spectra, distinctive retention time and comparison to authentic standards. Quantification was performed by integrating the peak areas with standard curves of authentic standards with the Waters millennium chromatography software. Lutein and β-carotene were relatively quantified at 450 nm and 270 nm respectively, by integrating their peak areas and calculating their percentage from total integrated peak areas. Tissues for chlorophyll determination were sampled as explained for carotenoid analysis. Chlorophyll extraction was performed in dimmed light to avoid possible photodegradation of chlorophyll. Chlorophyll was extracted by adding 5 mL of dimethyl sulfoxide (DMSO) to 0.5 g, vortexing and incubating in the dark at room temperature for 24 h. The extract was analyzed for absorbance in the wavelengths of 663 and 645nm using a Cary50Bio spectrophotometer (Varian). Chlorophyll concentration was calculated as described by Tadmor *et al*., (2010).

### Genotyping

DNA isolations were performed using the GenElute™ Plant Genomic Miniprep Kit (Sigma-Aldrich, St. Louis, MO). DNA quality and quantification were determined using a Nanodrop ND-1000 (Nanodrop Technologies, Wilmington, DE) Spectrophotometer, electrophoresis on agarose gel (1.0%) and Qubit^®^ dsDNA BR Assay Kit (Life Technologies, Eugene, OR).

### GBS analysis, SNP calling and map construction

(1) TAD×DUL RILs – DNA from 164 F_7_ individuals was processed by Novogene (Novogene Bioinformatics Institute, Beijing, China) for GBS analysis. 0.3∼0.6 μg of genomic DNA from each sample was digested with *ApeKI* restriction enzyme, based on the *in silico* evaluation results, and the obtained fragments were ligated with two barcoded adapters at each end of the digested fragment. Followed by several rounds of PCR amplification, all the samples were pooled and size-selected for the required fragments to complete the library construction. Samples where diluted to 1 ng/µl and the insert size was assessed using the Agilent® 2100 bioanalyzer; qPCR was performed to detect the effective concentration of each library. Libraries with a concentration higher than 2 nM were sequenced on an Illumina HiSeq 2000/2500 platform as 144 bp, paired-end reads and mapped to the *C. melo* reference genome DHL92 v3.5.1 (Garcia-Mas *et al.*, 2012; available at https://melonomics.net/fles/Genome/Melon_genome_v3.5.1/). Over 570 million reads were produced covering nearly 21% of the genome across more than 35 million tags at an average read depth of 9 reads per site. SNP calling was carried out using Broad Institute’s genome analysis toolkit (GATK) (McKenna *et al.*, 2010) resulting in 1,205,528 raw SNPs. Sites with a depth of less than three reads per site or more than 50 percent missing data were filtered out using TASSEL v5.2.43 (Bradbury *et al.*, 2007). Data was then imputed using full-sib families LD algorithm (Swarts *et al.*, 2014) followed by the removal of individuals with excess heterozygosity. The genotypic dataset was phased to ABH format consisting of 89,343 SNPs across 146 lines. Binning was performed using SNPbinner (Gonda *et al.*, 2018) with a minimum ratio between crosspoints set at 0.001 and minimum bin size of 1000 bp. Bin statistics and genetic distance were calculated using in-house script developed in python, based on the Kosambi mapping function (Kosambi, 1943). The final set included 2,853 recombination bins across 146 lines. Evaluation of genotypic data quality was done by accurately mapping flesh color to a 55Kb interval spanning the previously published *CmOr* gene (Melo3C05449) (Tzuri *et al.*, 2015).
(2) NA×DUL F_3:4_ – DNA from 140 F_3_ individuals was processed NRGene LTD (Nes Ziyyona, Israel) for Restriction-site-Associated DNA sequencing (RAD-seq) (Ríos *et al.*, 2017), SNP calling was performed following similar methods and the initial marker set included 43,975 SNPs across 140 individuals with an average depth of 16 reads per site. Further filtering and imputation were performed as described above and the final set for binning was composed of 19,015 SNPs across 134 individuals. Binning and genetic map construction were carried out using the same parameters as those used for the TAD×DUL population, yielding 1,321 bins across 134 individuals.
(3) GWAS180 - Genotyping of this diversity panel was performed using GBS, as described by Gur *et al.*, (2017). The final SNP set included 23,931 informative SNPs (at MAF>5%) across 177 accessions.
(4) The watermelon mapping population, NY0016×EMB, was genotyped by GBS. Library construction, sequencing, and SNP calling were performed at the Genomic Diversity Facility at Cornell University (Ithaca, NY) as described by Branham *et al.*, (2017). Sequences in this project were aligned to the Charleston Gray genome, version 1 (available at ftp://www.icugi.org/pub/genome/watermelon/WCG/v1/).

### Bulk Segregant Analysis by sequencing (BSA-Seq) of the watermelon population

DNA samples from 35 F_2_ plants (from NY0016×EMB cross) homozygote for the rind color trait (based on F_3_ family’s phenotypes) were prepared into two bulks (light rind: 19 F_2_ samples and dark rind: 16 F_2_ samples). These samples, as well as DNA samples of the parental lines (EMB and NY0016) were used for whole-genome resequencing performed at the DNA Services Center at the University of Illinois, Urbana-Champaign. Six shotgun genomic libraries were prepared with the Hyper Library construction kit from Kapa Biosystems (Roche) with no PCR amplification. The libraries were quantitated by qPCR and sequenced on one lane for 151 cycles from each end of the fragments on a HiSeq 4000 using a HiSeq 4000 sequencing kit version1. Fastq files were generated and demultiplexed with the bcl2fastq v2.17.1.14 Conversion Software (Illumina). Average output per library was 44 million reads of 150 bp. All raw reads were mapped to the Charleston Gray reference genome using the Burrows-Wheeler Aligner (BWA), producing analysis ready BAM files for variant discovery with Broad Institute’s Genome Analysis Toolkit (GATK). Homozygous SNPs between the two parents were extracted from the vcf file that was further filtered to total depth>20 reads per site. The read depth information for the homozygous SNPs in the ‘light’ and ‘dark’ pools was obtained to calculate the SNP-index (Takagi *et al.*, 2013). For each site we then calculated for each bulk the ratio of the number of ‘reference’ reads to the total number of reads, which represented the SNP index of that site. The difference between the SNP-index of two pools was calculated as ΔSNP-index. Sliding window method was used to perform the whole-genome scan and identify the trait locus confidence interval on chr9.

### Whole Genome re-Sequencing (WGS) of 25 representative diverse melon accessions

DNA of the 25 core accessions was shipped to the Genomic Diversity Facility at Cornell University (Ithaca, NY) for whole genome resequencing to an estimated 30X depth. *Validation of rare alleles at the APRR2 genes* – the causative variants at the different *APRR2* alleles in melon and watermelon, which were discovered based on NGS of genomic DNA, were confirmed on the parental lines and relevant segregants through Sanger sequencing of genomic DNA in melon and cDNA from mRNA that was extracted from fruits in watermelon.

### qRT-PCR analysis

Rind samples were peeled from fruits harvested throughout development, from anthesis to maturity, and immediately frozen in liquid nitrogen. Three fruits were sampled from each genotype at each developmental stage. 100-150 mg of frozen rind tissue per sample was used for RNA extraction using a Plant/Fungi Total RNA Purification Kit (NORGEN Biotek Corp., Canada). First-strand cDNA was synthesized using a cDNA Reverse Transcription Kit (Applied Biosystems, USA). The 10-μl qPCR volume included 1 μl of cDNA template, 0.2 μl of each primer (10 μM), 5 μl of Fast SYBR Green Master Mix (Applied Biosystems, USA), and RNase-free water to a final volume of 10 μl. qRT-PCR, with an annealing temperature of 60°C, was performed in triplicate on a 96-well plate in the Step-One Plus Real-Time PCR system (Applied Biosystems, USA). The melon *Cyclophilin A* gene (Melo3C013375) was used as a control to normalize the qRT-PCR values across different samples. Primers are listed in **Supplemental Table 5**.

### Data analysis

#### Trait mapping

Genome-wide linkage analysis for young fruit rind color was performed in TASSEL using a generalized linear model (GLM) in the bi-parental populations and confirmed using single-marker analysis in the JMP V13.1 software package (SAS institute, Cary, NC, USA). GWAS at the melon diversity collection was performed by a mixed linear model (MLM) analysis in TASSEL, using both the population structure (Q matrix) and relatedness (kinship (*k*) matrix) as covariates to control for population structure. Multiple comparisons correction to significance thresholds were performed using the FDR approach (Benjamini and Hochberg, 1995). All further statistical analyses (correlations and analyses of variance) were performed using the JMP V13.1 software package.

### Population structure, Kinship and LD analysis

Relatedness between the melon accessions in the diverse collection was estimated in TASSEL software v5.2.43 using the pairwise kinship matrix (k matrix) through the Centered IBS method. Linkage disequilibrium (LD) between intra-chromosomal pairs of sites was done on chromosome 4 using the full matrix option in TASSEL.

### Sequence analyses

Sequence alignments and comparison of *APRR2* alleles were performed using the BioEdit software package (Hall, 1999) and the integrative genomics viewer (IGV) package (Robinson *et al.*, 2011). Comparative analysis of haplotype diversity across 2,200 genes on melon chromosome 4 was performed following these steps: 1) A VCF file containing ∼4,000,000 high quality SNPs across the core set of 25 melon lines (MAF>0.1 and less than 10% missing data per SNP) was created based on alignments to the melon genome version 3.5.1. 2) The corresponding gene annotations file was used to create a subset of exonic SNPs on all annotated genes on chromosome 4. 3) The number of exonic-SNPs haplotypes per gene was calculated.

## Results

### GWAS of young fruit rind color in melon

Most melons can be visually classified into two distinct young fruit (∼10 days post anthesis) rind colors; dark or light green, reflecting qualitative variation in chlorophyll content. Light immature rind color was previously reported to display a recessive single-gene inheritance in a bi-parental segregating population (Burger *et al.*, 2006a). In the current study, young fruit rind color was visually scored on a previously described diverse melon collection composed of 177 accessions (Gur *et al.*, 2017). The collection was genotyped genome-wide with 23,931 informative, GBS-derived SNP markers, and was shown to be an effective resource for mapping simple traits in melon (Gur *et al.*, 2017). Here, we used a subset composed of 120 accessions with a clearly defined dark or light rind phenotype (Example in Figure 1a) for genome-wide association analysis. Accessions with prominent non-uniform rind color (stripes or dots) were excluded from this analysis. We also excluded *Charentais* lines, as their dominant grayish light rind is exceptional and phenotypically distinct from the common light rind in other melon types. While the dark and light phenotypes were distributed uniformly across the genetic variation and were represented in balanced proportions across this set (41% and 59%, respectively, Figure 1b), a genome-wide population-structure-corrected analysis did not result in any significant marker-trait association. This result has led to the assumption that while this highly heritable trait may show simple inheritance in a specific bi-allelic cross, it is possibly more complex and explained by multiple loci across a multi-allelic diverse collection.

**Figure 1:**
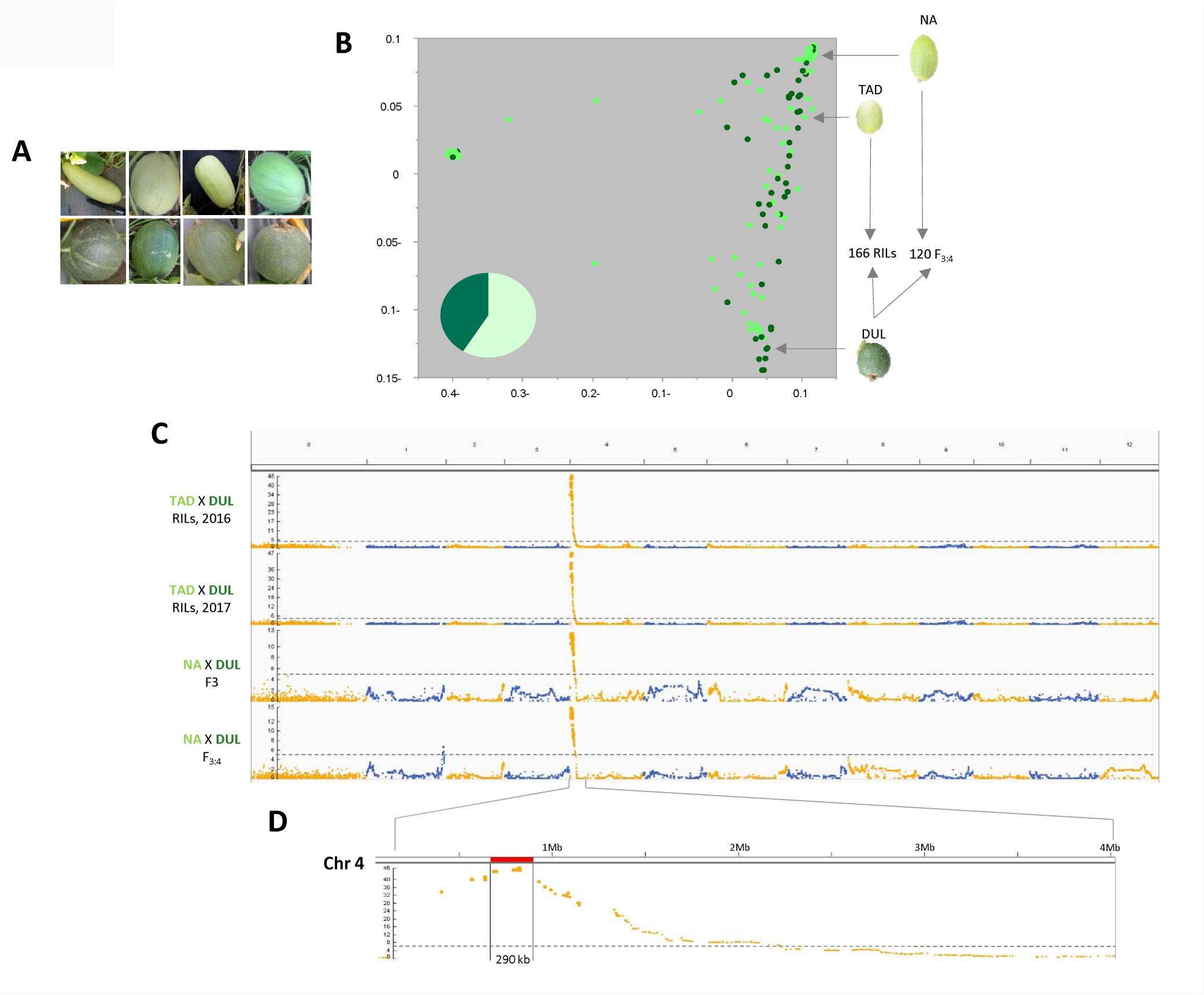
Characterization of variation and genetic mapping of young fruit rind color in melon. a) Example of accessions with light and dark rinds. b) Genetic PCA plot of 120 light and dark accessions from the diverse collection (Gur *et al.*, 2017). Dot color correspond to light and dark immature rind. Parental lines of the mapping populations are shown to the right of the plot (TAD: Tam Dew, NA: Noy-Amid). The pie chart on the left bottom corner summarizes the frequencies of young fruit rind colors. c) Manhattan plots for mapping of young fruit rind color across two populations over two seasons. d) Zoom in on chromosome 4. The 290Kb confidence interval is highlighted.

### Mapping and cloning of the young fruit light rind gene in melon

In order to further dissect this trait using a simpler genetic design, we analyzed two segregating bi-parental populations: the first is composed of 164 RILs (F_7_) from a cross between a light rind honeydew parent (Tam-Dew; TAD) and a dark rind *reticulatus* parent (Dulce; DUL, **Figure1b**). The second population is composed of 114 F_3:4_ families derived from a cross of DUL with another light rind accession, a yellow casaba *inodorous* melon (Noy-Amid; NA, Figure 1b). These segregating populations were visually phenotyped for young fruit rind color over two seasons and a consistent single gene (Mendelian) ratio was observed in dark:light phenotypes. The populations were then genotyped through GBS and 89,343 (TAD×DUL RILs) and 43,975 (NA×DUL F_3:4_) informative SNP markers were identified and used for mapping. Whole-genome linkage analysis using the four datasets (two populations over two growing seasons) resulted in the identification of a single highly significant consistent trait locus on chromosome 4 (Figure 1c). The common confidence interval for this trait locus spans a 290 Kb region (Chr4: 640-930 Kb, Figure 1d) on the melon reference genome (Garcia-Mas *et al.*, 2012; http://cucurbitgenomics.org/organism/3), as confirmed also through substitution mapping using recombinants within this interval in the TAD×DUL RILs population (Figure 2a). Annotation of the genomic sequence at this interval revealed 33 putative genes (**Sup. Table 1**) including a strong candidate, Melo3C003375, which is annotated as an *ARABIDOPSIS PSEUDO-RESPONSE REGULATOR2-LIKE* (*APRR2)* gene, the melon homolog of a recently reported causative gene of the recessive white rind (*w*) mutation in cucumber (Liu *et al.*, 2016).

**Figure 2:**
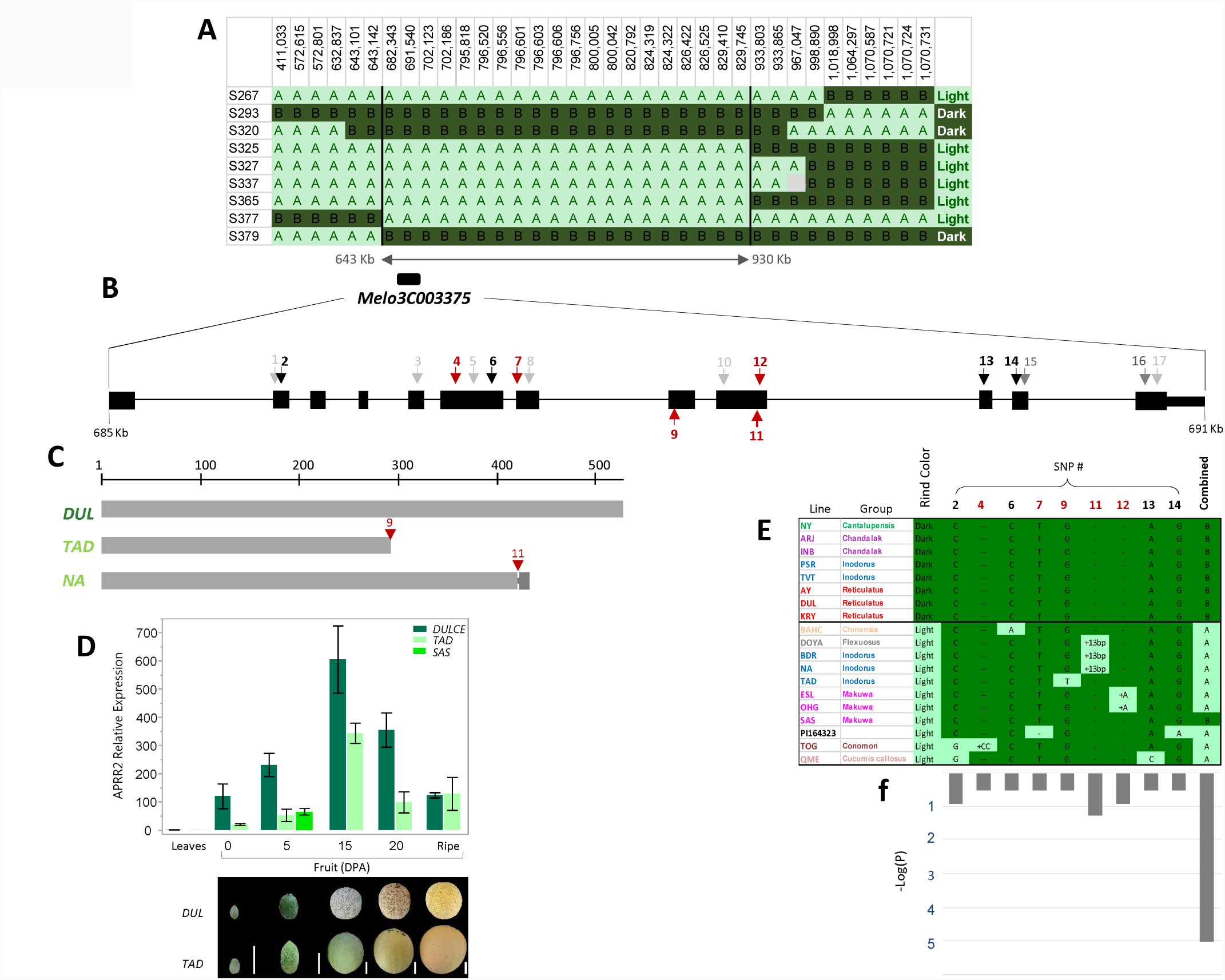
Fine mapping and candidate gene characterization. a) Substitution mapping at the TAD×DUL RILs population. Nine recombinant RILs at the QTL interval are shown. Marker bins physical positions (bp) are indicated on the top of each column. Light and dark colors correspond to parental alleles (TAD and DUL, respectively). RILs young fruit rind color phenotypes are shown on the right column. Trait mapping interval is bounded with thick vertical lines. Position of the candidate gene Melo3C003375 is shown. b) Melo3C003375 gene structure and exonic sequence variants. Black boxes represent exons. Light-gray arrows represent synonymous SNPs. Dark-gray arrows are non-synonymous SNPs that did not show a distinct allelic state between dark and light accessions (15, 16). Black arrows are SNPs causing a single amino-acid change, and red arrows are polymorphisms (SNPs or InDels) causing major change in protein (frame-shift, stop codon). c) Predicted protein size of the dark (DUL; Dulce) and light rind parents (TAD; Tam-Dew, NA; Noy-Amid). d) Expression pattern of the *CmAPRR2* gene through fruit development, and comparison between parental lines. DPA: days post anthesis. e) Table of non-synonymous allelic variants in Melo3C003375 that distinguish between light and dark phenotypes across 19 diverse melon lines. Accessions are colored by their horticultural group. f) Association tests of independent causative variants and combined ‘functional variant’ with rind color, across the core set.

We then compared the *CmAPRR2* gene (Melo3C003375) sequence between the parental lines of the mapping populations. Genomic and mRNA sequencing revealed multiple polymorphisms, including two different exonic polymorphisms causing independent stop codons in each of the light rind parents compared to the common dark parent (DUL). G to T substitution in exon 8 in TAD compared to DUL lead to a premature stop-codon and a predicted aberrant protein of 292 amino-acids (AA) compared to the normal 527 AA protein of DUL (Figure 2b-c**, Sup Figure 1**). A 13-bp insertion in exon 9 of NA result in a frame-shift leading to a different premature stop-codon in this line and a predicted protein of 430 AA (Figure *2*c**, Sup Figure 1**). Furthermore, we crossed TAD and NA with each other and with DUL (as a reference testcross) and phenotyped the F_1_s for young fruit rind color. Both testcrosses with DUL resulted, as expected, in a dark rind in the F_1_. However, the F_1_ of TAD×NA had a clear light rind and confirmed the allelic nature of these recessive phenotypes (Figure 3a). This further corroborates that these independent predicted causative mutations in the *CmAPRR2* gene are indeed allelic.

**Figure 3:**
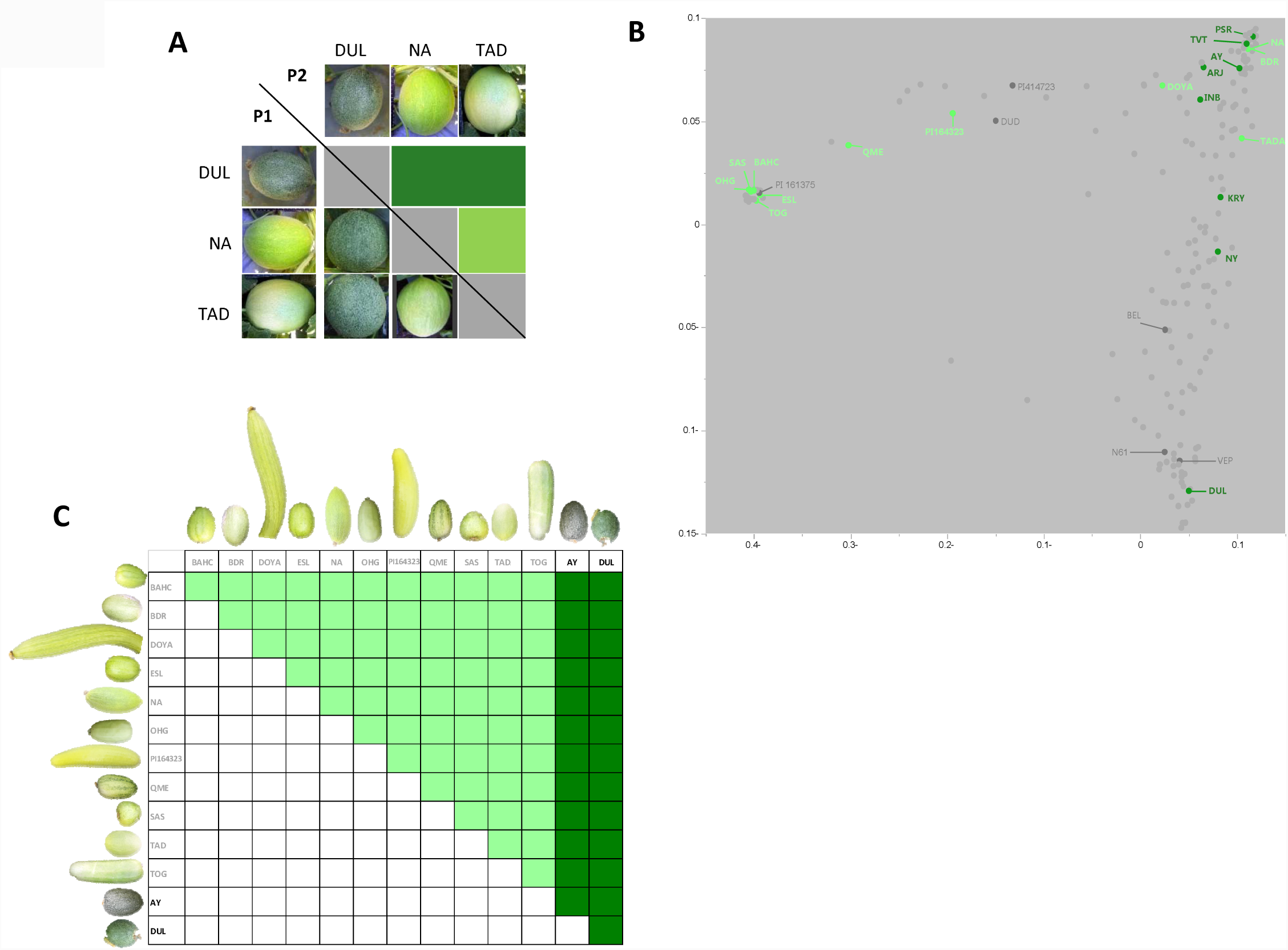
Allelism tests for light rind accessions. a) Allelism test for the mapping populations light rind parental lines (TAD and NA). b) Genetic PCA plot with 25 selected founders highlighted by rind color. In gray are lines with non-distinct rind color. c) half-diallele allelism tests across 11 light rind accessions. Two dark lines were used as reference testers.

### Expression pattern of the CmAPRR2 gene in melon fruit

Fruits from the light (TAD) and dark (DUL) parental lines were sampled during development from anthesis to maturity and mRNA levels of the *CmAPRR2* gene were analyzed by qRT-PCR. We show here that *CmAPRR2* has higher expression level in fruit comparted to leaves (Figure 2d), as shown also in cucumber (Liu *et al.*, 2016) and pepper (Brand *et al.*, 2014), and in agreement with the Melonet-DB gene expression atlas (Yano *et al.*, 2018). In accordance with these studies, we also show that the *CmAPRR2* peak expression in fruit rind occur around 15 days post anthesis (DPA), before the initiation of ripening and color change. Comparison between the parental lines of the mapping population also showed significantly lower levels of *CmAPRR2* expression in light rind fruits throughout fruit development. Both light and dark lines have reduced *CmAPRR2* expression levels at the mature fruit stage and were not significantly different from each other at that stage (Figure 2d).

### Multiple independent causative mutations in the CmAPRR2 gene across melon diversity

In light of the conflict between the clear identification of a single causative gene through linkage mapping in two different bi-parental crosses on one-hand, and the absence of any significant genome-wide signal from the GWAS analysis on the other hand, we re-sequenced and compared the genomic sequence of the *CmAPRR2* gene across a core panel of 25 diverse melon lines. This core panel was selected to represent the different groups and overall diversity in our collection as described previously (Figure 3b) (Gur *et al.*, 2017). Nineteen lines from the panel, showing a clear dark or light young fruit rind phenotype, were used for the sequence comparison. Seventeen SNPs and InDels within exons in the *CmAPRR2* gene were identified across this panel (Figure 2b). Eight of these SNPs were either synonymous or did not show a distinct allelic state between dark and light accessions (in gray). The remaining nine polymorphisms are independently inherited (not in LD with each other) and display low frequency alleles (0.05-0.15) which are unique to the light rind accessions (Figure 2e). Four of these polymorphisms (2, 6, 13 and 14) are SNPs that change a single amino acid (in black). Three are InDels (4, 7 and 12) that cause frame-shifts leading to major modification in predicted protein sequence (in red), and the remaining two (9 and 11) are the causative polymorphisms described above, leading to premature stop codons as in TAD and NA. These five major polymorphisms (4, 7, 9, 11 and 12) explain the light rind phenotype in eight of the eleven light rind accessions in the core panel. The non-synonymous polymorphisms 2, 6 and 13 are potentially causative of the light rind phenotype in BAHC and QME. The only light rind accession that could not be explained by non-synonymous variation within the *CmAPRR2* coding sequence is SAS. However, the low mRNA expression of *CmAPRR2* in young fruit (5 DPA) rinds of SAS, which was similar to the expression level in TAD, and significantly lower compared to DUL (Figure 2d), suggest expression level variation as a possible causative element for the light rind phenotype in this line. To test whether all these ‘light’ accessions are indeed allelic and caused by different mutations in the *CmAPRR2* gene, we performed allelism tests where all the ‘light’ accessions (n=11) were intercrossed and the resulting 55 F_1_s were phenotypically evaluated for young fruit rind color. As a reference, these ‘light’ accessions were crossed with two ‘dark’ testers (DUL and Ananas Yoqne’am; AY). Figure 3c shows that all 55 ‘light’×’light’ F_1_ hybrids displayed light immature fruit rinds, while all 22 ‘light’×’dark’ testcrosses displayed dark rinds. These results confirm the allelism between the 11 ‘light’ lines, including the light rind phenotype of SAS.

Combined interpretation of the sequence variation and allelism tests across this representative core panel indicate that most of the young-fruit rind color variation can be explained by multiple independent polymorphisms related to the *CmAPRR2* gene. Extrapolation of the results from this core set suggests that each of these causative variants is most likely also present at low allele frequency, across the wider diversity panel, leading to the non-significant associations observed in our GBS-based GWAS experiment. It is worth noting that these independent mutations are not in LD with each other, leading to the high haplotype diversity in this gene. In a comparative analysis of haplotype diversity based on exonic SNPs across 2,200 genes on melon chromosome 4, we found that *CmAPRR2* is indeed the second most diverse gene, irrespective of number of SNPs and transcript length (**Sup Figure 2** and methods). Analysis of the LD pattern in the genomic region surrounding the *CmAPRR2* locus, confirmed the low LD between SNPs in this region (**Sup Figure 3**). The fact that these allelic polymorphisms are not in LD with each other allow us aggregate them into a theoretical unified functional polymorphism, resulting in increased frequency of aberrant *CmAPRR2* allele (0.52, Figure 2e right column). This analysis, in turn, produces a significant association between the *CmAPRR2* gene and the light rind trait (Figure 2f).

### Mapping and cloning of the light rind color gene in watermelon

In parallel with melon, we also studied the genetics of rind color in watermelon. The main difference is that in watermelon, due to its non-climacteric fruit ripening, chlorophylls are the main rind pigments also during fruit maturity and therefore light and dark green were visually scored on mature fruits. Our light rind source in this study was an heirloom accession named NY0016 (Tadmor *et al.*, 2005) that was crossed on different lines, varying in their rind color, to produce F_1_s and F_2_s. All the F_1_ hybrids had dark rind fruits (irrespective of the stripes pattern in the ‘dark’ parents), proving the recessive nature of the light rind phenotype of NY0016. All F_2_ populations were phenotyped for rind color and a consistent 3:1 Mendelian ratio was observed for dark and light rinds, respectively (Figure 4a, **Sup Table 2**). The cross between the light rind accession (NY0016) and a striped (dark) parent (Early Moon Beam; EMB) was selected for linkage mapping and this population was advanced to F_3_ to perform F_2:3_ analysis (Figure 4b). GBS of the F_2_ population (N= 87) resulted in a final high-quality set of 3,160 filtered SNPs (Branham *et al.*, 2017) that was used for genetic mapping of the light rind phenotype. Seven to ten fruits per F_3_ family were visually scored and each family was classified into a defined category: fixed for light rind, fixed for dark rind or segregating for rind color (**Sup. Figure 4**). The observed 1:2:1 frequencies of light, segregating and dark across the F_3_ families supported a single gene inheritance for this trait (Figure 4b). Whole-genome linkage analysis resulted in the identification of a single significant trait locus on chromosome 9 (R_2_=0.62, P=2.9×10_-18_, Figure 4c). The confidence interval for this locus spanned 1.7 Mb with 80 predicted genes on the watermelon reference genome (Charleston Gray, http://cucurbitgenomics.org/organism/4). In order to narrow down this genomic interval, we performed mapping-by-sequencing of DNA bulks (BSA-seq) from 35 selected F_2_ segregants that were homozygous for light or dark rind (on the basis of fixed F_3_ phenotypes). The ∼30X whole-genome re-sequencing resulted in the identification of 400,000 high quality SNPs differentiating between parental lines. Comparison of allele frequencies between the light and dark bulks across the 400,000 SNPs (ΔSNP-index analysis) confirmed the trait locus on chromosome 9 and allowed us to narrow the genomic confidence interval to a 900Kb region with 30 predicted genes (Figure 4d**, Sup Table 3**). Review of the list of genes within the confidence interval revealed a strong candidate, CICG09G012330, the watermelon homolog *ClAPRR2* gene, highly similar to the causative melon (Melo3C003375) and cucumber (Csa3G904140.3 (Liu *et al.*, 2016)) genes. Comparison of the genomic sequence of CICG09G012330 between the mapping population parental lines revealed several SNPs, none of them within exons. The only putative causative SNP at that point was at intron 6/exon 7 junction (Figure 4e). Parents and segregants mRNA sequence comparison revealed an alternative splicing in the intron6/exon7 junction, leading to a 16 bp deletion at the mRNA of the light rind parent and corresponding segregants carrying the ‘light’ allele. This 16 bp deletion, which created a frame shift leading to a premature stop codon and a predicted aberrant protein, is most likely causative for the light rind phenotype of NY0016 (Figure 4f, g).

**Figure 4:**
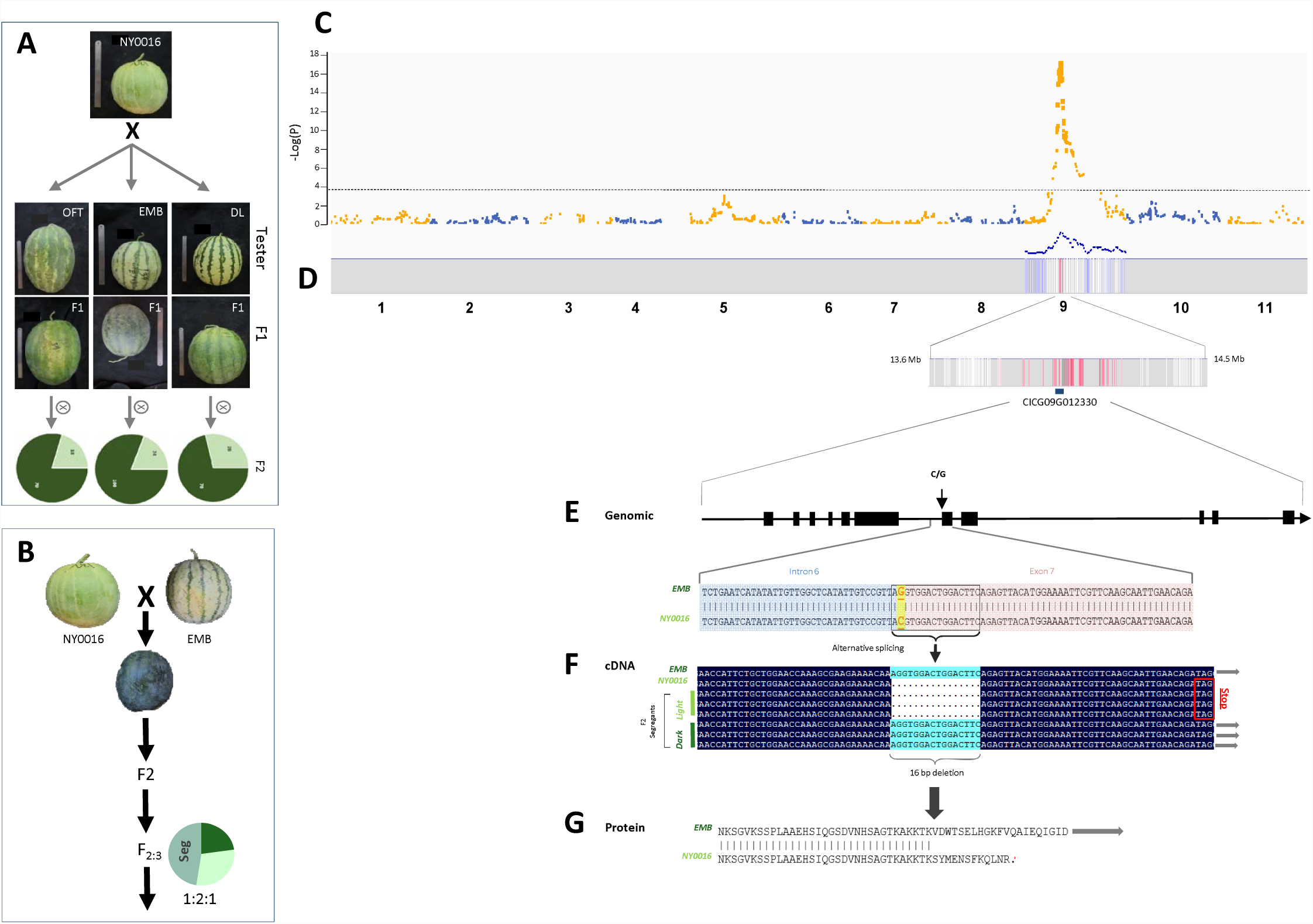
Mapping and cloning of the light rind color gene in watermelon. a) Three testcrosses and F_2_ segregation for rind color. OFT: Orange Flesh Tender sweet, EMB: Early Moon Beam, DL: Dixie Lee. b) Parents of the mapping population and rind color frequency distribution in the F_2:3_ population. c) Manhattan plot of whole-genome linkage analysis in the F_2:3_ population using 3,160 GBS-derived SNPs. d) BSA-Seq results across 35 fixed F_2:3_ families. Association significance (δSNP-index analysis) is expressed using blue-to-red color scale. Position of the watermelon *APRR2* gene (CICG09G012330) is shown. e) *ClAPRR2* (CICG09G012330) annotated gene structure. Exons are represented as black boxes. Parental genomic sequence alignment and SNP (C/G) at the intron 6 exon 7 junction. f) Comparison of mRNA sequence of parental lines (EMB and NY0016) and three segregants from each rind color group. Stop codon downstream to the 16 bp indel is shown. g) Alignment of parental lines translated protein sequence around the InDel site.

### Allelic variation in the CmAPRR2 gene is associated with mature fruit rind and flesh pigmentation in melon

Earlier analyses of rinds from TAD, DUL and selected F_3_ families from their cross, demonstrated the correlation between young fruit chlorophyll content and mature fruit carotenoids (**Sup. Figure 5**). To test whether the *CmAPRR2* gene is also associated with mature fruit rind pigmentation in melon, we analyzed mature fruits from the TAD×DUL RILs population for rind color and carotenoids content. We harvested 10 mature fruits per line, external images of the fruits were taken for color scoring and the rinds were sampled for carotenoids profiling. Color intensity variation at the young (green) stage in this population is qualitative and could be visually classified into two distinct classes (dark or light green) on a single-fruit basis. However, at the mature stage any effect of the *CmAPRR2* gene on rind color is visually of a quantitative nature (Figure 5a), as shown also in tomato (Pan *et al.*, 2013). Rind netting, which segregates in this population, further masked rind color and complicated visual scoring and carotenoids quantification. For the parental lines, TAD, with the light green rind at young fruit stage, has a cream-yellowish rind at maturity and DUL, with the dark green rind at young stage, has an orange rind masked by dense rind netting (Figure 2d). Visual observation of standardized external images of selected mildly-netted mature fruits, representing both alleles in the *CmAPRR2* gene, suggested a possible effect of this gene on rind color intensity, such that on average the ‘dark’ allele is associated with deeper orange color (Figure 5a). We confirmed this effect through analysis of carotenoids content in rinds of 50 selected mildly netted segregants (25 RILs carrying each allele in the *CmAPRR2* gene). The ‘light’ allele was significantly associated with more than 10 fold reduction in total carotenoids in the rind, which is consistent with the reduced chlorophyll levels observed in young fruits of this group (Figure 5b**, Sup Figure 5**). Allelic variation in this gene is explaining 37% of the variation in lutein content (P=2×10^-6^), 34% of the variation in β-carotene content (P=8×10^-6^), and 37% of total carotenoids (P=3×10^-6^) in fruit rind in this population.

**Figure 5:**
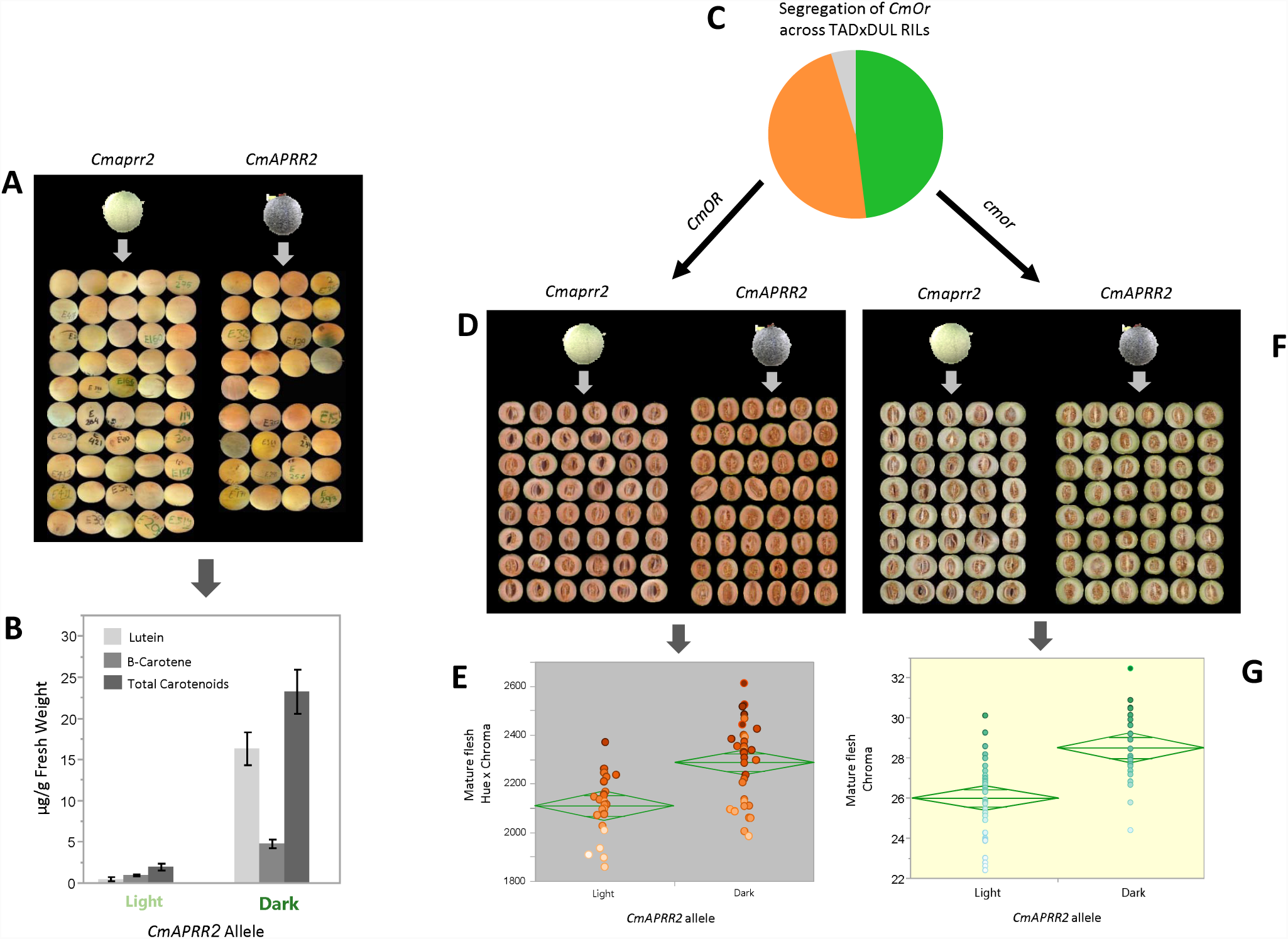
Association of allelic variation in the *CmAPRR2* gene with mature fruit rind and flesh pigmentation in TAD×DUL RILs. a) External images of fruits from RILs with dark and light genotype in the *CmAPRR2* gene. b) Analysis of carotenoids in mature fruit rinds. c) Segregation of orange and green flesh controlled by the *CmOr* gene, across 166 RILs. d) Representative scans of orange fruits segregating for dark and light alleles at the *CmAPRR2* gene. e) Analysis of *CmAPRR2* allelic effect on flesh color of mature orange (*CmOR/CmOR*) fruits. f) Representative scans of green fruits segregating for dark and light alleles in the *CmAPRR2* gene. g) Analysis of *CmAPRR2* allelic effect on flesh color of mature green (*cmor/cmor*) fruits.

To examine whether the effect of the *CmAPRR2* gene extends also to mature fruit flesh color, we phenotyped the TAD×DUL RILs for flesh color intensity using longitudinal fruit section scanning and quantitative image analysis (n=145 lines x 10 fruits per line). This population is segregating for the main flesh color gene in melon, *CmOr*, discriminating between orange and non-orange flesh (Tzuri *et al.*, 2015) and it is therefore composed of orange flesh lines (48%) and green flesh lines (49%), the remaining 3% of the lines are segregating due to residual heterozygosity (Figure 5c).

In order to test the association of the *CmAPRR2* gene with flesh color, we analyzed the orange and green fruits separately. A significant association between the *CmAPRR2* allelic segregation and color intensity was found in the orange-flesh group, which accumulated β-carotene as the main flesh pigment (R^2^=0.25, P=4.7×10^-5^, Figure 5d-e) and as expected, the ‘dark’ allele was associated with stronger pigmentation and higher predicted β-carotene content (R^2^=0.62, P=0.0053, see Materials and methods and **Sup. Figure 6**). Significant association of this gene with flesh color was found also in the green flesh group, which accumulated chlorophyll as the main flesh pigment (R^2^=0.31, P=2×10^-6^) as the ‘dark’ allele was significantly associated with higher green Chroma (Figure 5f-g) reflecting higher chlorophyll content.

### Expression level of the CmAPRR2 gene is associated with mature flesh color in melon

In order to test whether the expression level of the *CmAPRR2* gene is associated with mature fruit pigmentation, we analyzed RNA-seq data and mature fruit flesh carotenoids on a different RILs population derived from a cross between DUL and an Indian phut snapmelon (Momordica group), PI414723 (hereafter called 414) (Galpaz *et al.*, 2018). While 414 has a spotted rind (and not a clear light or dark phenotype), we assume based on testcrosses with some of the light rind accessions that, as DUL, it also carry a ‘dark’ allele of the *CmAPRR2*. This assumption is supported by the fact that it does not show any of the predicted ‘light’ non-synonymous polymorphisms found at the *CmAPRR2* gene (Figure 2 b, e). DUL and 414 are genetically and phenotypically distant and differ in their mature fruit flesh color and carotenoids content (Figure 6a). DUL has dark orange flesh while 414 has light (salmon-colored) orange flesh, and accordingly the RILs population segregates for these traits (Harel-Beja *et al.*, 2010; Galpaz *et al.*, 2018). RNA-Seq was previously performed on mature fruit flesh of 96 RILs from this cross (Freilich *et al.*, 2015) and now allowed us to execute a genome-wide eQTL analysis for the *CmAPRR2* gene (Melo3C003375). A single, highly significant, cis-eQTL was mapped to chromosome 4 and defined by 270 Kb interval flanking this locus (Galpaz *et al.*, 2018, Figure 6b-c). This result confirmed the heritable variation in *CmAPRR2* expression level in this population and that the expression of this gene is mostly regulated by cis-acting sequence variants. We then tested the correlation between *CmAPRR2* expression level and flesh β-carotene content across the 96 RILs and found a significant positive correlation (R=0.38, P=0.0008, Figure 6d). Since both parental lines of this population carry the ‘dark’ allele based on the coding sequence of the *CmAPRR2* gene, this result provides a quantitative support for the possible relationship between expression level of this gene and pigment accumulation in melon. It is important to note that this population is segregating at additional QTLs that affect carotenoids content in mature fruit flesh, including a major QTL on chromosome 8, which was recently mapped to a candidate gene level (Diaz *et al.*, 2011; Galpaz *et al.*, 2018). This variation further masked the specific effect of *CmAPRR2* expression level on flesh carotenoids content. We also assume that the observed correlation is an underestimation, as gene expression in this experiment was measured on mature fruits whereas the peak of expression of the *CmAPRR2* gene is much earlier before fruit ripening (∼15 DPA).

**Figure 6:**
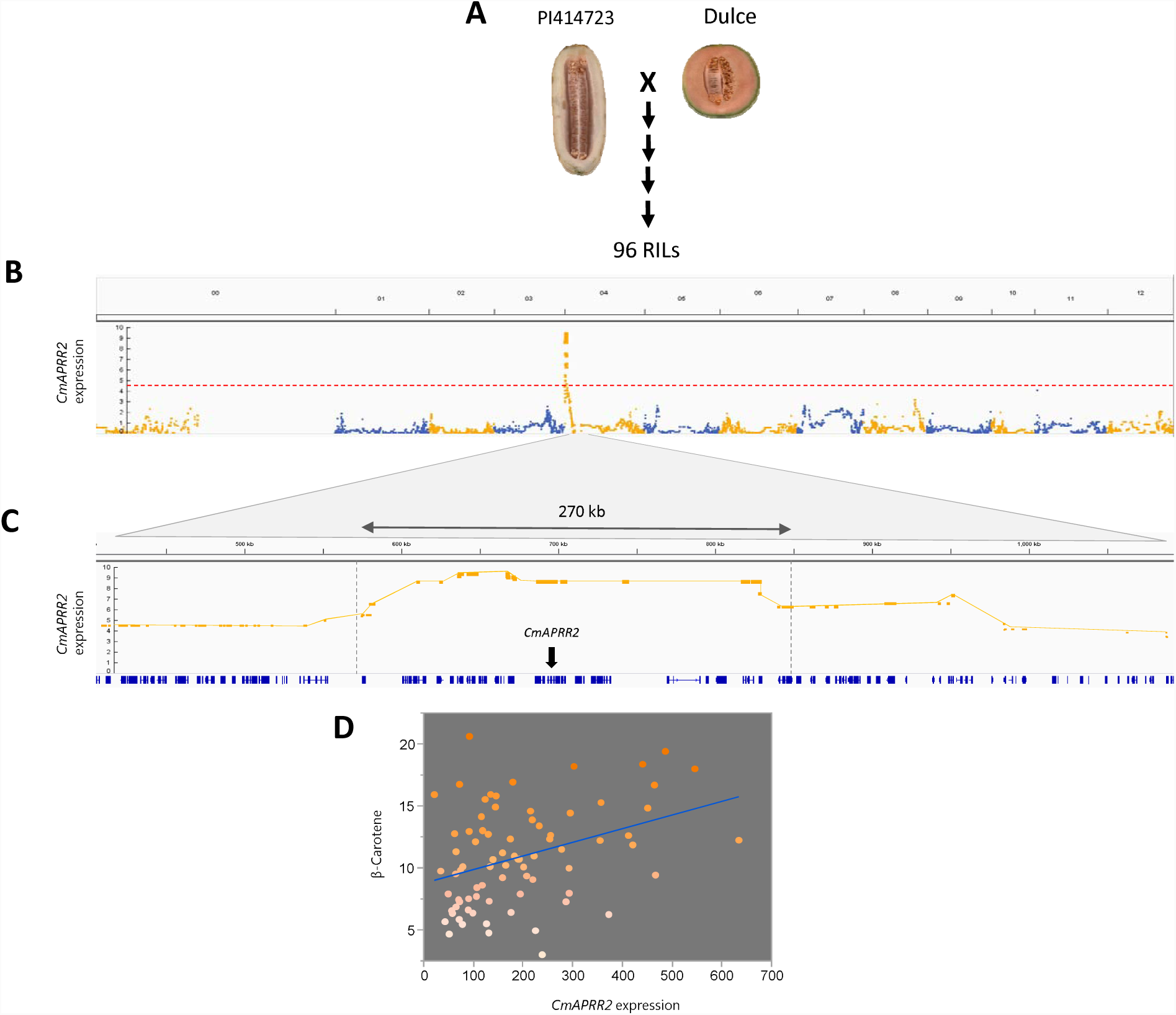
*CmAPRR2* (Melo3C00375) is differentially expressed, and correlated with β-carotene content in the 414×DUL RILs. a) Population parents (DUL and 414). b) Manhattan plot for whole-genome eQTL mapping of *CmAPRR2* (Melo3C003375) expression (RPKM) in mature fruit flesh. c) Zoom in on cis-eQTL spanning Melo3C003375 on chr.4. d) Correlation between Melo3C00375 expression (RPKM) and β-carotene content (µg/g fresh weight) in the fruit flesh.

### Expression of CmAPRR2 in melon is correlated with plastid-development related genes

To characterize co-expression patterns associated with the *CmAPRR2* gene, we calculated the correlations between the expression of Melo3C003375 and all annotated melon genes (n=27,557), using RNA-Seq data from mature fruits of the 414×DUL RILs population (n=96) (Freilich *et al.*, 2015). Fourteen thousand genes expressed in mature fruit flesh were used for this correlation analysis. Gene ontology (GO) enrichment analysis was performed on 200 genes that had the strongest correlations with Melo3C003375 (R>0.43, FDR adjusted P<0.001). The four most significant functional groups that were enriched are related to photosynthesis, light reaction and plastid organization and the fifteen most enriched components are related to plastids and chloroplasts (**Sup Table 4**). These results support the predicted involvement of the *CmAPRR2* gene in the regulation of chloroplasts and chromoplasts development.

## Discussion

### Color variation in immature fruit rind in melon

External fruit color is an important attribute in melons as it is a key factor defining consumers’ preference. Melon rind color transforms during fruit development and ripening, mainly by shifting from the green rind of immature fruits, where chlorophyll is the main pigment, to variable rind colors composed of different combinations of chlorophylls, carotenoids and flavonoids (Tadmor *et al.*, 2010). Inheritance of external color of immature fruit has been previously described in two different studies: The white color of immature fruits was reported by Kubicki, (1962) to be dominant to green immature fruits and controlled by a single gene named *Wi* (Dogimont, 2011). Burger *et al.*, (2006*a*) described a recessive gene for light immature exterior color in a cross between an American muskmelon (*Reticulatus* group) and an American honeydew-type melon, (*Inodorous* group). Recently, a major gene for external color of immature fruit was mapped in a cross between “Védrantais”, a *Charentais* line from the *Cantalupensis* group, and “Piel de Sapo”, from the *Inodorous* group (Pereira *et al.*, 2018). The dominant light rind from the “Védrantais” parent, that most likely correspond to the *Wi* gene, was mapped to ∼1.6 Mb interval on chromosome 7 (Pereira *et al.*, 2018). These results confirm our observation that the dominant light grayish rind of *Charentais* accessions is phenotypically and genetically distinct and controlled by a different gene from the one we identified in the current study, which correspond to the recessive gene described by Burger *et al.*, (2006*a*).

### APRR2-like transcription factors are key regulators of fruit pigmentation

The results of the current study support the pivotal role of *APRR2-like* genes in regulation of pigmentation in fruits. Pan *et al.* (2013) showed that over expression of an *APRR2* gene in tomato resulted in increased chlorophyll content in immature fruits and higher carotenoids level in ripe tomatoes. Both effects resulted from an increase in plastid number. They also provided evidence for association between null mutation in an *APRR2* gene and external fruit color intensity in green peppers. These results are complementary to reports on the role of a related transcription factors group, GLKs, which were shown to be associated with levels of chlorophyll and carotenoids in Arabidopsis, tomato and pepper (Waters *et al.*, 2008, 2009; powell *et al.*, 2012; Brand *et al.*, 2014). Recently, Liu *et al.*, (2016) reported that an *APRR2* gene is causative for the white rind (*w*) mutation in cucumber, expressed as reduced chloroplast density and chlorophyll content in young cucumber fruits. In all these crop plant species (tomato, pepper and cucumber), there is also a correlation between expression levels of either *APRR2* or *GLK* genes and pigments intensity. In the current study, we performed high-resolution NGS-based mapping in segregating populations and found that null mutations in the *CmAPRR2* and *ClAPRR2* genes are associated with light rind color in melon and watermelon, respectively. We also showed that expression of the *CmAPRR2* gene is correlated with pigment intensity in melon (Figures 2d, 6d) and that, as in pepper (Brand *et al.*, 2014) and cucumber (Liu *et al.*, 2016), these transcription factors show their strongest expression in fruit and reach their peak expression around 10-20 DPA and before fruit ripening. Our results expand the extent of experimental data that demonstrate the conserved function of *APRR2*-like genes in regulating fruit pigmentation as shown by the comparable expression profiles and analogous phenotypes associated with variation in these genes.

### CmAPRR2 is associated with pigment accumulation across fruit developmental stages and tissues

Variation in ripe fruit color is substantially wider compared to the variation in the immature stage. While in the immature stage color variation mostly reflect chlorophyll concentrations, in the mature stage biosynthetic pathways of additional pigments (i.e. carotenoids, flavonoids) are involved, leading to extended complexity of the genetic architecture. This complexity is also expressed by the independent genetic control of flesh and rind colors in melon as best demonstrated by the ability to combine different rind and flesh colors through classical breeding. Rind and flesh color QTLs were mapped in multiple melon populations (Monforte *et al.*, 2004; Cuevas *et al.*, 2008, 2009; Harel-Beja *et al.*, 2010; Galpaz *et al.*, 2018; Pereira *et al.*, 2018) and a few causative color genes were identified (Feder *et al.*, 2015; Tzuri *et al.*, 2015). However, so far, a common regulator that affects pigmentation throughout the different developmental stages and fruit tissues (rind and flesh) has not been described in melon. In the current study, we showed that the *CmAPRR2* gene is such a key regulator, associated with pigments concentrations during the course of fruit development and across fruit tissues (Figures 1, 5). Furthermore, the mapping population in this study (TAD×DUL RILs), which independently segregated for both *CmOr* and *CmAPRR2* genes, allowed us to show that the *CmAPRR2* effect is also independent of the type of pigment accumulated in the flesh, as it was associated with variation in both chlorophyll and carotenoids concentrations (Figure 5c-g). We also showed here, using a different segregating population (414×DUL RILs) that was previously subjected to mature fruit RNA-Seq and carotenoid analysis (Freilich *et al.*, 2015; Galpaz *et al.*, 2018), that the cis-regulated *CmAPRR2* expression variation is correlated with flesh β-carotene content in mature fruits (Figure 6). Since both parental lines of this population carry a predicted ‘dark’ allele based on the coding sequence of the *CmAPRR2* gene, we assume that this experiment provided another piece of evidence for the relationship between expression level of *CmAPRR2* and flesh pigments content. The proposed involvement of this transcription factor in regulation of plastid development in fruits (Pan *et al.*, 2013; Liu *et al.*, 2016) support the broad effect of *CmAPRR2* that was described here.

### Multi-allelic nature of the CmAPRR2 gene in melon

The genetic architecture that describes a phenotypic trait is strongly dependent on the type and structure of the germplasm used for the study. For quantitative polygenic traits, QTL segregation and detection will not necessarily overlap across different mapping populations. The same can apply to simple traits, where independent mutations in different genes, which are involved in a common biological process, lead to the same discrete phenotype. While bi-parental populations will draw only part of the picture in such cases, diverse collections or multi-parental segregating populations are more effective in comprehensively characterizing this architecture. In the current study we tried to genetically characterize the light immature rind phenotype in melon using a diverse collection, assuming it is under a simple genetic control as previously described (Burger *et al.*, 2006a). In GWAS, lack of detection power can result from low heritability, low frequency of the phenotype under investigation, strong confounding effect of population structure or insufficient markers density (Korte and Farlow, 2013). While none of these factors seemed to apply in our case (Gur *et al.*, 2017 and Figure 1), we did not obtain any significant GWA signal, which led to the intuitive assumption that multiple genes are associated with the light immature rind phenotype in our collection. The identification of two independent allelic nonsense mutations in the *CmAPRR2* gene, through linkage analyses (Figures 1, **2**), indicated that we might be looking at a different scenario. Through complementary resequencing of a diverse core panel and comprehensive allelism testing (Figures 2, 3) we were able to demonstrate that this trait is a unique case of simple genetic architecture. On the functional level, it seems to be controlled by a single gene that segregates in a Mendelian manner in bi-parental crosses, but the multi-allelic pattern at the *CmAPRR2* gene drove reduced power through GWAS, which masked this simplicity and created the observed contradiction between the different mapping strategies. These results provide a thought-provoking example for another possible inherent complexity that can arise in GWAS - independent low-frequency causative variants within a common gene. In the current scenario, even whole-genome deep resequencing of the GWAS panel, which would target each of the variants in the *CmAPRR2* gene, would not necessarily resolve the lack of detection power, as each of these independent variants remain at low frequency. Genetic mapping studies are rapidly shifting towards sequencing-based genotyping and in most cases marker density is no longer a bottleneck in GWAS (Yano *et al.*, 2016; Misra *et al.*, 2017). A key challenge remains in prioritizing GWAS signals and improving weak signals obtained from low frequency causative variants (Lee and Lee, 2018). The availability of whole-genome assemblies and corresponding protein-coding gene annotations, alongside additional layers of information, such as expression profiles from RNA-Seq experiments, now facilitate the integration of multiple data layers to improve GWAS results (Shim *et al.*, 2017; Lee and Lee, 2018; Schaefer *et al.*, 2018). Our example of the *CmAPRR2* gene suggests that adding functional annotation prediction to GWAS SNPs and treating predicted genes as integral functional units could potentially be used as an informative layer that can boost the signal of causative weak associations.

In summary, we have identified the *CmAPRR2* gene as a common regulator of fruit pigmentation in melon and watermelon. The conserved and broad effect of this gene across species, fruit tissues, developmental stages and different types of pigment accumulated, suggest its potential as a useful target for carotenoids bio-fortification of cucurbits and other fruits.

## Supplementary data

**Supplementary Figure 1**: cDNA sequence comparison between the mapping population parental lines DUL, TAD and NA.

**Supplementary Figure 2:** Number of annotated transcript haplotypes vs number of SNPs per bp (a) and transcript length (b) across 2,200 genes on chromosome 4. The *CmAPRR2* gene is highlighted in red.

**Supplementary Figure 3**: *CmAPRR2* gene is located in a low LD region. a) LD (R^2^) heat map by physical position in a 400Kb window surrounding the *CmAPRR2* gene. b) Sliding-window trend line of pairwise SNP LD across 1M bp interval surrounding the *CmAPRR2* gene on chr4.

**Supplementary Figure 4:** Segregation and scoring of rind color of 87 F_3_ families from the NY0016×EMB cross. a) Examples of the three phenotypic rind color classes in the population. b) Segregation of the color classes across F_3_ families.

**Supplementary Figure 5**: Analysis of rind chlorophyll and carotenoids during fruit development on TAD, DUL and selected F_3_ families from their cross. a) Analysis of rind chlorophyll content during fruit development in TAD and DUL. b) Comparison of rind chlorophyll content between light and dark F_3_ families at 20 DAA. c) Comparison of rind total carotenoids content between light and dark F_3_ families at maturity (40 DAA). d) Correlation between chlorophyll content at 20 DAA and carotenoids at 40 DAA across the selected F3 families and parental lines.

**Supplementary Figure 6:** Prediction of flesh β-carotene based on fruit-section image analyses in the TAD×DUL RILs. a) Analysis of flesh color (Chroma) on fruit sections scans of ∼700 orange fruits in the TAD×DUL RILs. b) Calibration curve for the relation between flesh Chroma and β-carotene concentration as measured across 73 diverse orange flesh accessions from our GWAS panel. Logarithmic equation representing the best fit is shown. c) Calculation of predicted β-carotene in the TAD×DUL RILs based on Chroma values from (a) that were placed in the equation described in (b). Analysis of predicted β-carotene concentrations across the TAD×DUL RILs. Comparison between the ‘light’ and ‘dark’ alleles in the *APRR2* gene.

**Supplementary Table 1:** Melon young fruit light rind QTL interval: annotations and positions of genes.

**Supplementary Table 2:** Segregation of light rind in four F_2_ watermelon populations.

**Supplementary Table 3:** Light rind QTL interval in watermelon: annotations and positions of genes.

**Supplementary Table 4:** Gene Ontology (GO) enrichment analysis for 200 genes correlated with expression of the *APRR2* gene (Melo3C003375) in melon fruit.

**Supplementary Table 5:** list of primers used for RT-qPCR.

## Acknowledgments

We wish to thank the Newe-Yaar farm team for technical assistance in setting the field trials. We thank Jeff Glaubitz and the team at the Genomic Diversity Facility at Cornell University and Alvaro G. Hernandez and his team at the DNA Services center at the University of Illinois for the sequencing services for this research. Funding for this research was provided by the Israeli Ministry of Agriculture Chief Scientist grant no. 20-01-0141, by the United States-Israel Binational Agricultural Research and Development Fund (BARD) grant no. IS-4911-16.

