## Supplementary figures and images for "Multi-allelic *APRR2* Gene is Associated with Fruit Pigment Accumulation in Melon and Watermelon"

### Supplemental Figures

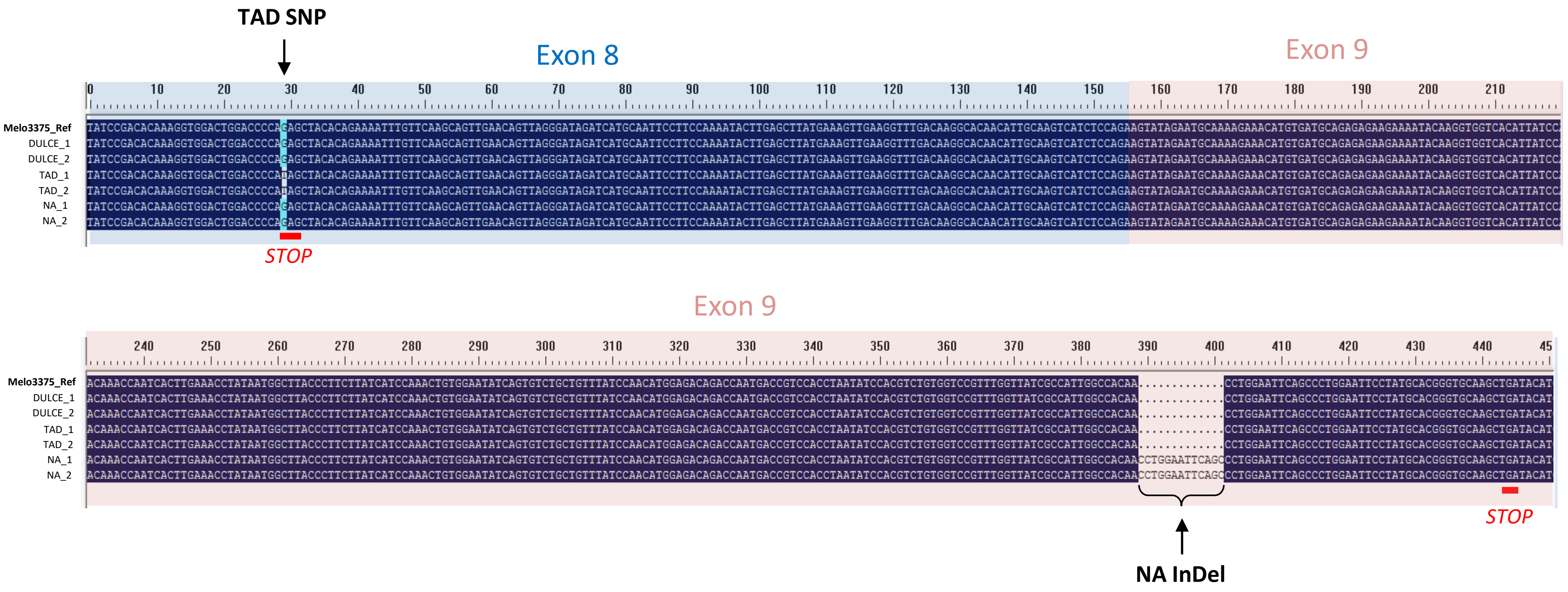

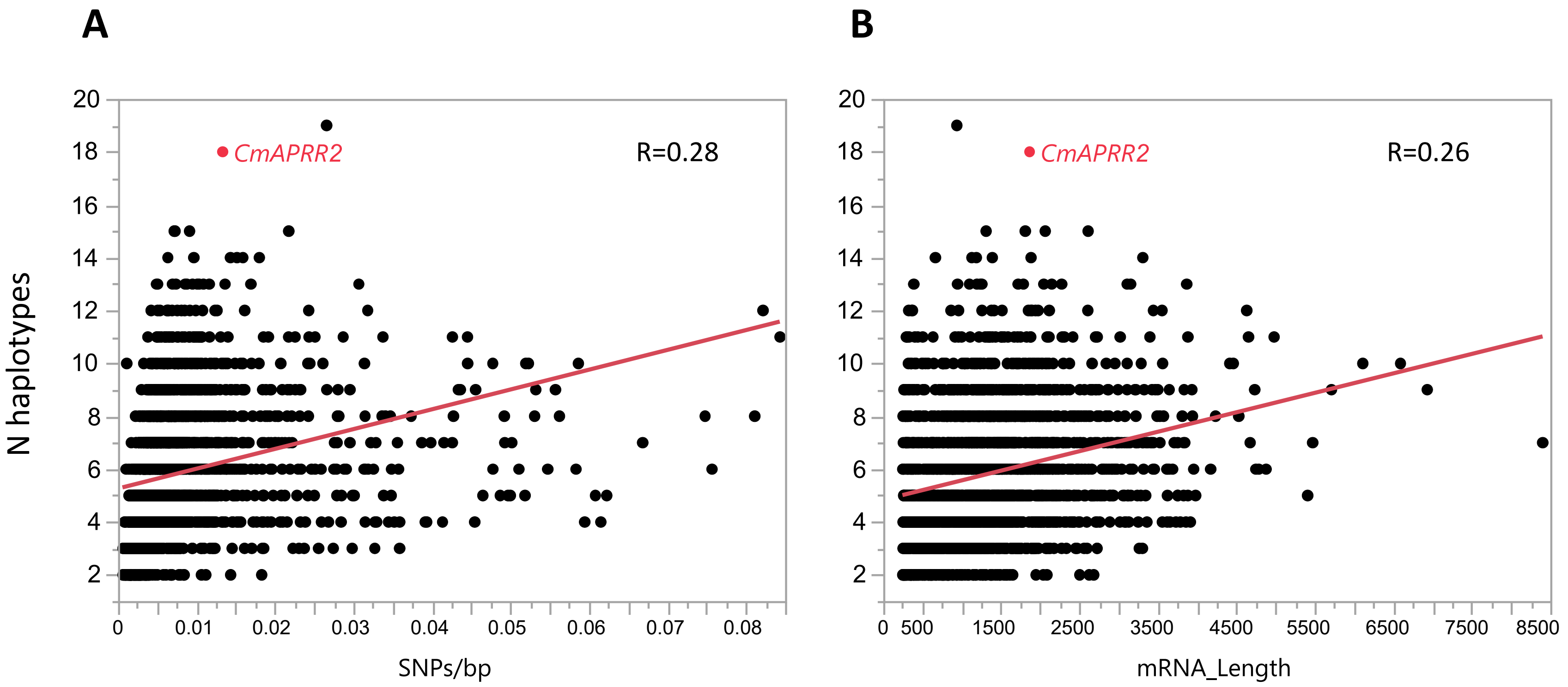

Sup Figure 3

A

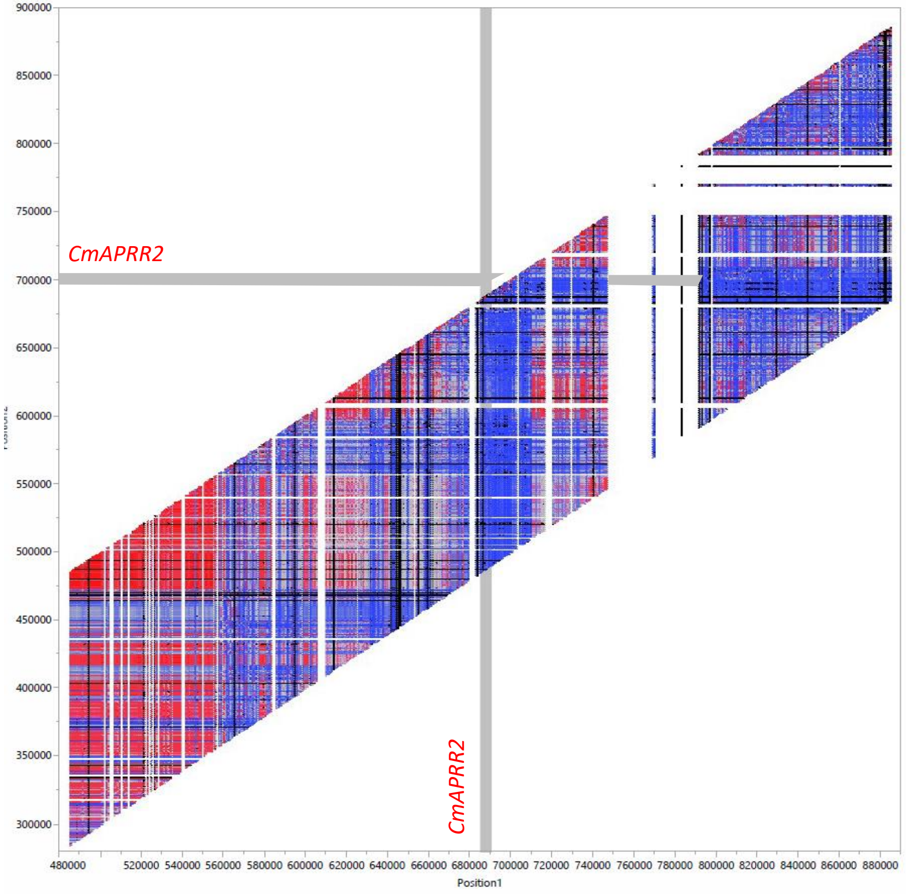


B

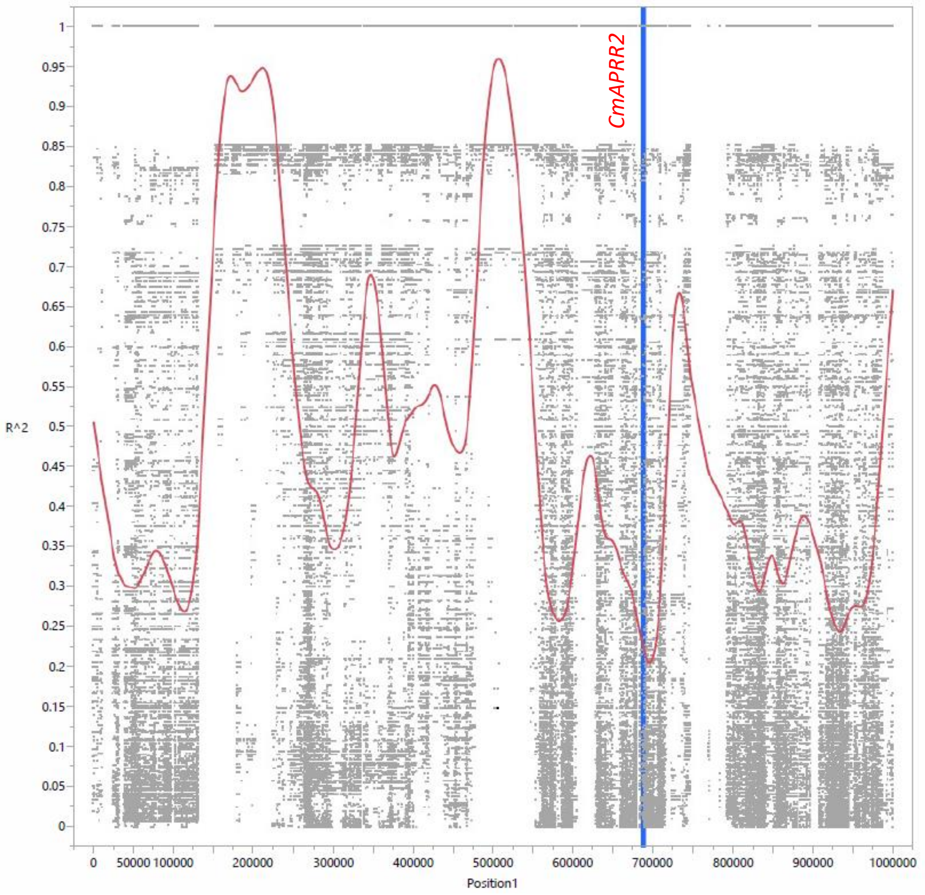

**Sup Figure 4**

A

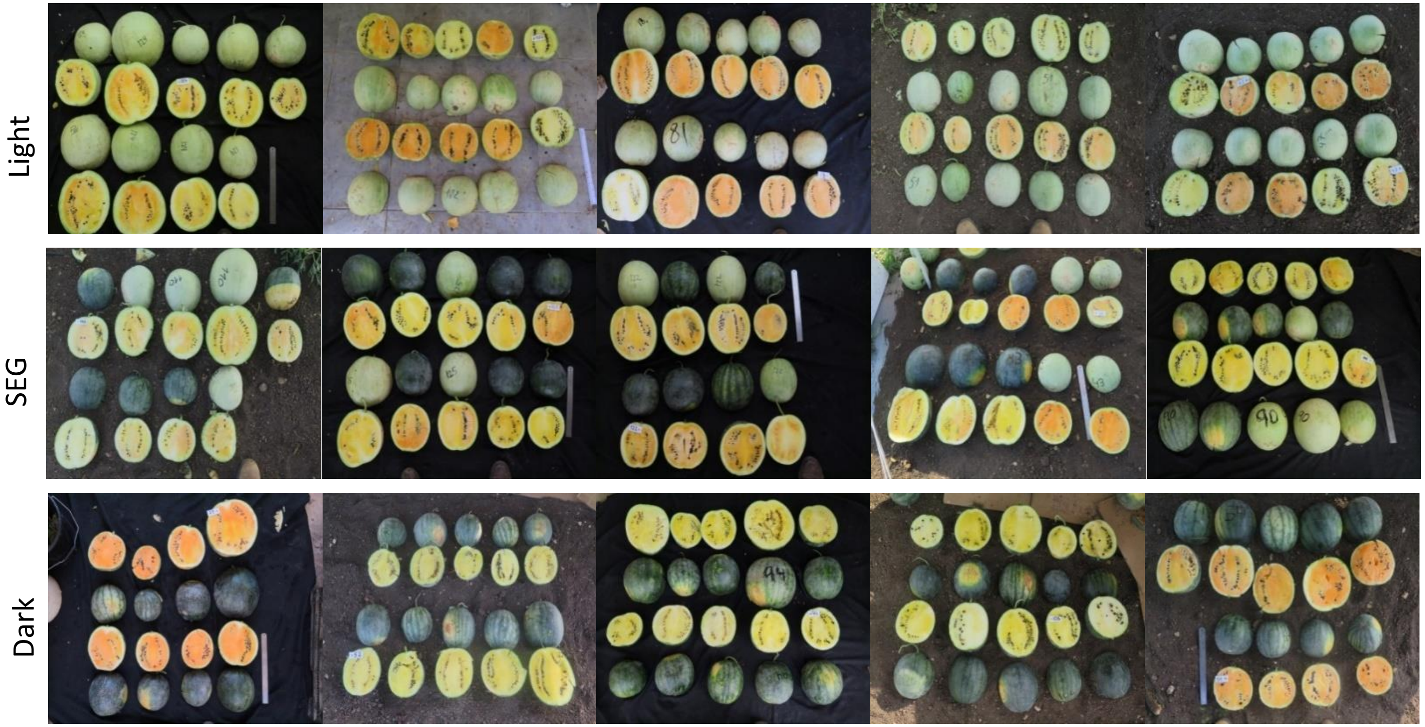


B

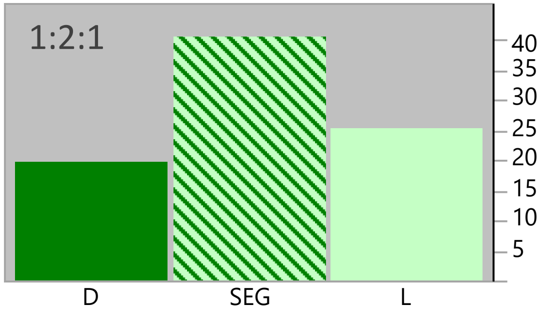

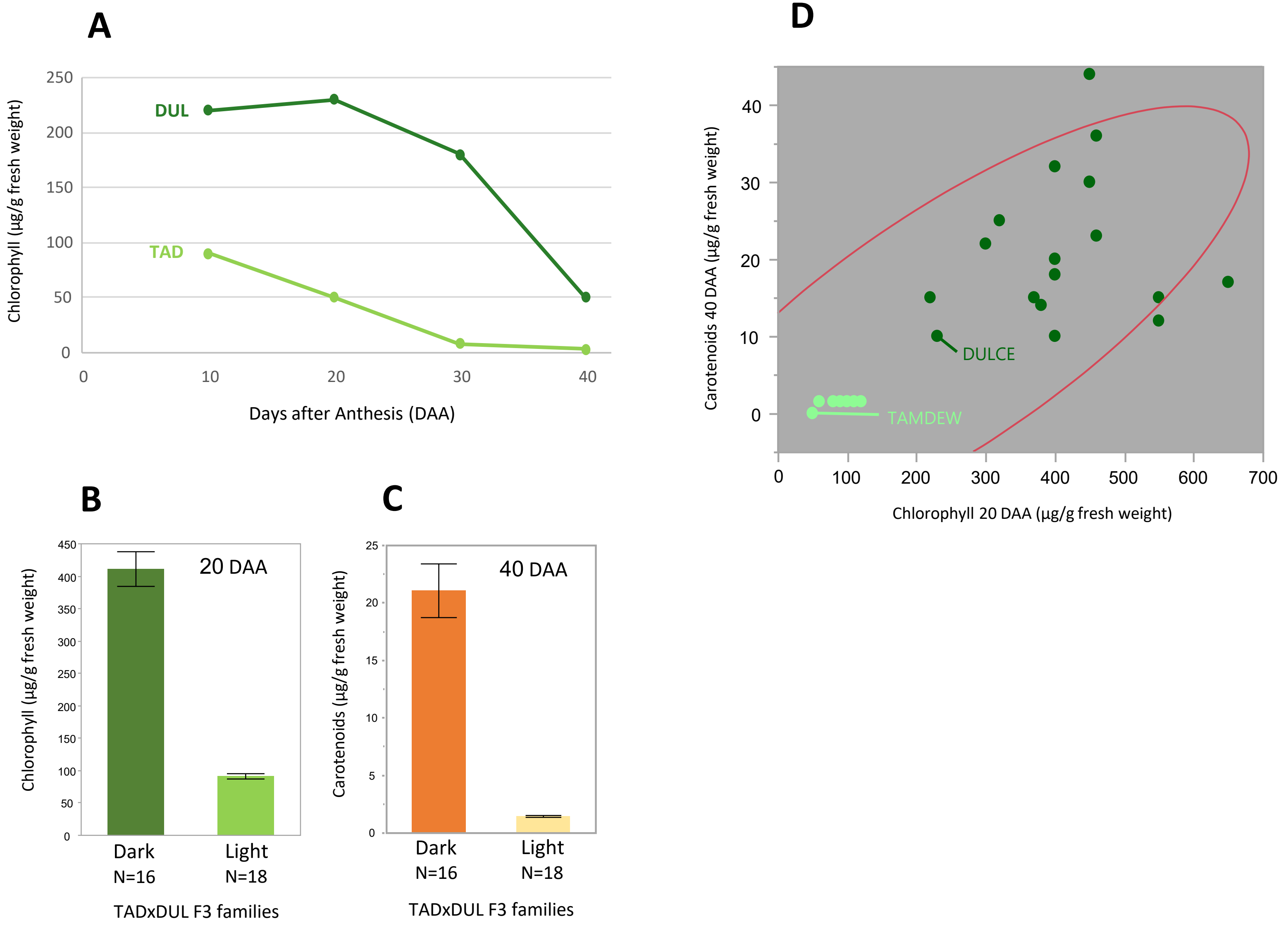

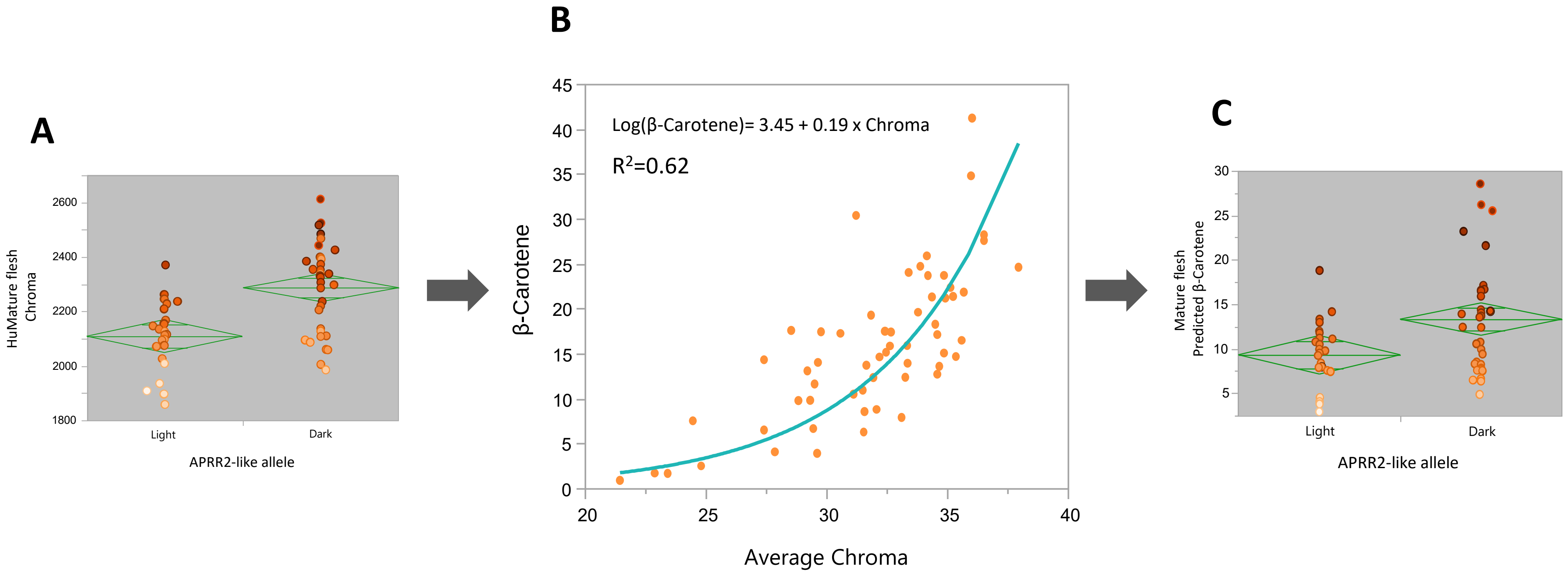
